# Evolutionary selection of proteins with two folds

**DOI:** 10.1101/2023.01.18.524637

**Authors:** Joseph W. Schafer, Lauren L. Porter

**Affiliations:** National Library of Medicine, National Center for Biotechnology Information, National Institutes of Health, Bethesda, MD 20894, USA; National Heart, Lung, and Blood Institute, Biochemistry and Biophysics Center, National Institutes of Health, Bethesda, MD 20892, USA

**Keywords:** Protein Fold Switching, Coevolutionary Analysis, Protein Evolution, Protein Structure Prediction, Protein Folding, Conformational Diversity

## Abstract

Although most globular proteins fold into a single stable structure^1^, an increasing number have been shown to remodel their secondary and tertiary structures in response to cellular stimuli^2^. State-of-the-art algorithms^3-5^ predict that these fold-switching proteins assume only one stable structure^6,7^, missing their functionally critical alternative folds. Why these algorithms predict a single fold is unclear, but all of them infer protein structure from coevolved amino acid pairs. Here, we hypothesize that coevolutionary signatures are being missed. Suspecting that over-represented single-fold sequences may be masking these signatures, we developed an approach to search both highly diverse protein superfamilies–composed of single-fold and fold-switching variants–and protein subfamilies with more fold-switching variants. This approach successfully revealed coevolution of amino acid pairs uniquely corresponding to both conformations of 56/58 fold-switching proteins from distinct families. Then, using a set of coevolved amino acid pairs predicted by our approach, we successfully biased AlphaFold2^5^ to predict two experimentally consistent conformations of a candidate protein with unsolved structure. The discovery of widespread dual-fold coevolution indicates that fold-switching sequences have been preserved by natural selection, implying that their functionalities provide evolutionary advantage and paving the way for predictions of diverse protein structures from single sequences.

## Introduction

Though machine learning methods have recently revolutionized protein structure prediction^3,5,8^, some systems remain a challenge^9-12^. For example, fold-switching proteins, also known as metamorphic proteins^13,14^, transition between two sets of stable secondary and tertiary structure^2,15^. These structural transitions modulate protein functions involved in suppressing human innate immunity during SARS-CoV-2 infection^16^, regulating the expression of bacterial virulence genes^17^, maintaining the cycle of the cyanobacterial circadian clock^18,19^, and more^20^. In spite of their biological importance, AlphaFold2 predicts only one conformation for 92% of these dual-folding proteins, and 30% of the predicted conformations were likely not the lowest energy state^21^. Other structure prediction algorithms, such as trRosetta^22^ and EVCouplings^4^, also systematically failed to predict experimentally validated fold switching in the universally conserved NusG family of transcription factors^7^.

Most state-of-the-art protein structure prediction algorithms, including all just mentioned, infer folding information from evolutionary conservation patterns. Very early studies of protein structure^23^ recognized that amino acid pairs with similar selection rates, also known as coevolved residue pairs, tend to be in direct contact^24,25^. These coevolved contacts can greatly constrain the number of possible conformations that computational methods must sample to predict a protein’s fold^26^, motivating the development of increasingly sophisticated methods that infer amino acid coevolution^27-33^. Multiple sequence alignments (MSAs), collections of sequences likely homologous to the sequence of interest, are the inputs to most of these methods. Typically, the accuracy of inferred coevolved residue pairs increases with MSA depth^34^, though recent deep learning-based methods can make accurate inferences from shallow MSAs^5,35^.

Some previous work hints that amino acid contacts unique to both conformations of fold-switching proteins may have coevolved. This work demonstrates that alternative protein conformations can be detected in subfamily-specific MSAs^7,36^. For example, we recently identified fold-switching proteins within the universally conserved NusG transcription factor family by leveraging structural information derived from MSAs from protein superfamilies (deep MSAs) and protein subfamilies (shallow MSAs with sequences similar to a target of interest)^7^. Furthermore, Dishman and colleagues found that several reconstructed ancestors of the fold-switching chemokine XCL1 switch folds, from which they concluded that XCL1 fold switching was evolutionarily selected^37^. These studies, though suggestive, focus on a couple of specific systems and infer fold switching from experimental observation of a few variants (XCL1) or inconsistent secondary structure predictions (NusG). Weak coevolutionary couplings of a fold-switching NusG have also been predicted, though the couplings had high proportions of noise: 38% at best^38^.

To robustly search for amino acid contacts unique to both conformations of fold-switching proteins, we applied unsupervised learning techniques to both superfamily and subfamily-specific MSAs of all 98 known fold switchers with two distinct experimentally determined structures^2^. One technique identifies coevolution of amino acid pairs using Markov Random Fields (MRFs). The MRF construction offers several advantages: (i) it converges to a global minimum as MSA depth increases, (ii) it can generate reasonable predictions from fairly shallow MSAs, and (iii) the MRF formalism accounts for noncausal correlations that arise when two residues interact with a third but not with one another^39-41^. Among the numerous MRF-based methods^4,42-44^, we selected GREMLIN (Generative REgularized ModeLs of proteINs) because of its superior performance^38,45^. The second technique, MSA transformer, infers coevolved amino acid pairs using a language model that focuses on both evolutionary patterns of amino acids within an MSA (column-wise attention) and properties of the individual sequences (row-wise attention), often with better accuracy than GREMLIN for single-fold proteins^35^.

We gauge the success of these methods by quantifying the overlap between predicted and experimentally determined residue-residue contacts from both folds. These comparisons are easily visualized with contact maps, which display amino acid pairs either measured or predicted to be proximal (heavy atom distance ≤8Å^45^). Though typical contact maps are symmetric about the diagonal, those used here are asymmetric to maximize information content. For example, the large light gray circles in the upper triangular portion of **Figure 1** represent contacts unique to the experimentally determined monomeric fold of KaiB, while the black circles in the lower triangular portion represent contacts unique to KaiB’s experimentally determined tetrameric fold. Contacts common to both experimentally determined folds are shown in medium gray on both sides of the diagonal. Where appropriate, interchain contacts are represented by smaller circles using the same color scheme. Predicted contacts, though not shown in Figure 1, are smaller and teal. Correct predictions are opaque circles; incorrect predictions are translucent diamonds.

**Figure 1:**
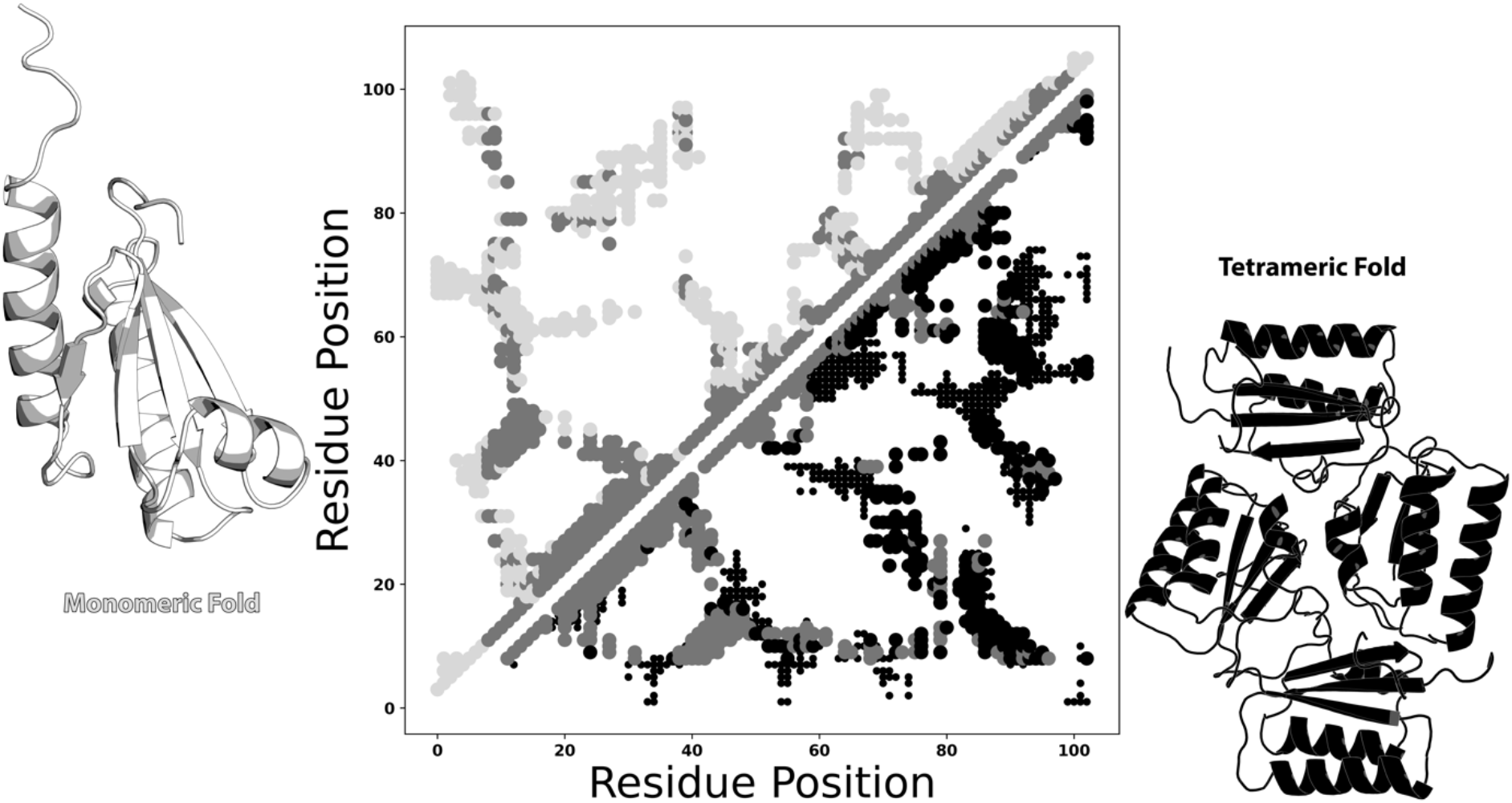
Example of a dual fold contact map from experimentally determined structures. KaiB monomeric/tetrameric contacts within 8Å are shown in the upper/lower triangles of the contact map in in light gray/black. Contacts common to both folds are shown in dark gray. Interchain contacts within 10Å are shown as smaller circles in their respective colors. Monomeric/tetrameric contacts were calculated from PDBs 5JYT/4KSO. Protein structures were generated with PyMOL^46^. Plots in all figures were generated with Matplotlib^47^.

## Results

### Approach to identify dual-fold contacts

**Figure 2** depicts our workflow to search for dual-fold coevolution (**Methods**). The query sequence, which corresponds to two distinct experimentally determined structures, is used to generate a deep MSA. This MSA is filtered to create successively shallower MSAs with sequences increasingly identical to the query (**Figure 2a**). These increasingly subfamily-specific MSAs are intended to reveal coevolutionary couplings from alternative conformations, as they did with RfaH, a fold-switching NusG protein^48^ whose ground state α-helical conformation can be only in subfamily-specific MSAs^7^. Accordingly, coevolutionary analysis is performed on each MSA using GREMLIN and MSA Transformer (**Figure 2b**). Predictions from both methods run on these nested MSAs are combined and superimposed on a single contact map (**Figure 2c**). Finally, these predictions are filtered by density-based scanning^49^ to remove noise (**Figure 2d**).

**Figure 2:**
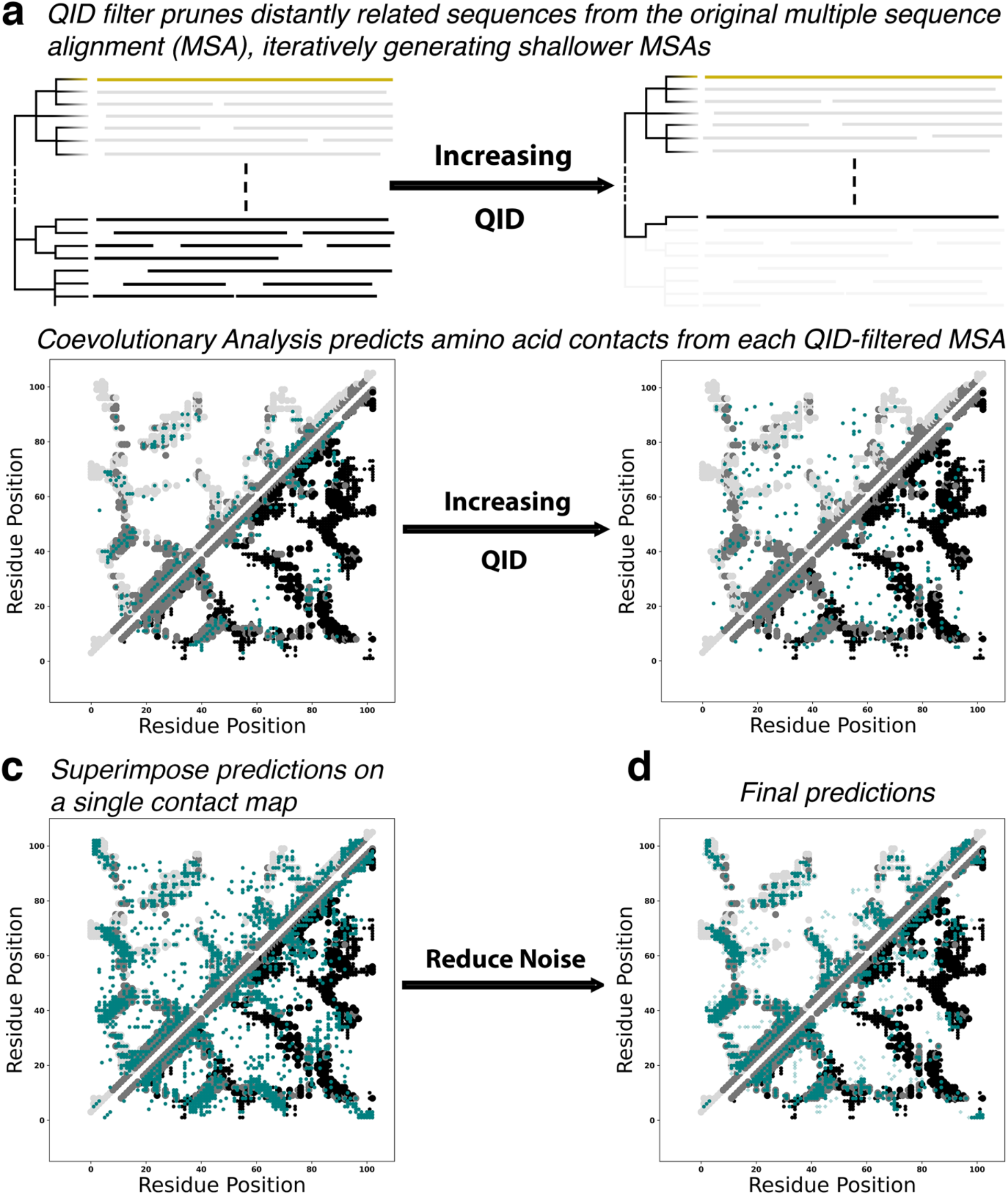
Graphical depiction of the workflow designed for applying coevolutionary analysis to fold-switching proteins. a) An MSA suitable for coevolutionary analysis is pruned using a Query Identity (QID) filter, removing distantly related sequences from the dataset and generating subfamily-specific MSAs. b) Each MSA (original + all pruned) is used as input for coevolutionary analysis. c) Predictions from all MSAs are superimposed on a single contact map. d) A clustering algorithm filters noise, leaving dense clusters of predicted amino acid contacts. Contacts unique to the dominant/alternative folds are light gray/black; common contacts are light gray; experimentally consistent predictions are teal circles; incorrect predictions (noise) are translucent teal diamonds.

Contacts are categorized as follows. Dominant fold: unique contacts corresponding to the experimentally determined structure that overlaps most with predicted contacts from the deepest MSA (light gray contacts in **Figure 2b-d**); Alternative fold: unique contacts corresponding to the other experimentally determined structure (black contacts in **Figure 2b-d**); Common: predicted contacts overlapping with experimentally determined contacts shared by both folds (gray contacts on both diagonals in **Figure 2b-d**); Noise: predicted contacts that do not overlap with any experimentally determined contacts (readily visible in **Figure 2c**). Though we call it noise, these experimentally inconsistent contacts could result from yet another alternative—but experimentally uncharacterized—conformation.

#### Evolutionary selection of dual-fold proteins

We applied our approach to all known fold-switching proteins, 91 single sequences with two distinctly folded experimentally determined structures^2^. These proteins are found in all kingdoms of life and represent >80 distinct fold families (**Extended Data Table 1**). Although efforts were made to generate the deepest possible MSA for each fold-switching sequence (**Methods**), the depths of 32 MSAs were too shallow for downstream analysis (<5*length of query sequence^45^) and one displayed severe artifacting after analysis (**Extended Data Table 1**). Thus, the rest of our approach was applied only to the remaining 58 fold-switching sequences with sufficiently deep MSAs (**Extended Data Table 1, Extended Data Figures 1**). Conformations with more contacts predicted in the deep MSA are denoted “dominant”, and those with fewer predicted contacts, “alternative”. This terminology holds no biophysical significance: 33% of “dominant” conformations do not correspond to the lowest energy states (**Extended Data Table 2**).

Our approach predicted substantially more correct contacts than coevolutionary analysis run deep superfamily MSAs alone, the standard approach^34^. The number of correctly predicted contacts increased for 56/58 proteins, with mean/median increases of 46%/41% (**Figure 3a)**. Predicted amino acid contacts uniquely corresponding to the 58 alternative conformations were highly enhanced, with mean/median increases of 98%/69% (**Figures 3b**). Noise was amplified substantially less than correctly predicted alternative contacts, with mean/median increases of 50%/30% (**Extended Data Figure 2a**). Prior to density-based filtering, mean/median noise was amplified by 377%/277%, demonstrating that, on average, 85% of the extra noise accrued from subfamily MSAs is sparsely distributed (**Extended Data Figure 2b**).

**Figure 3.**
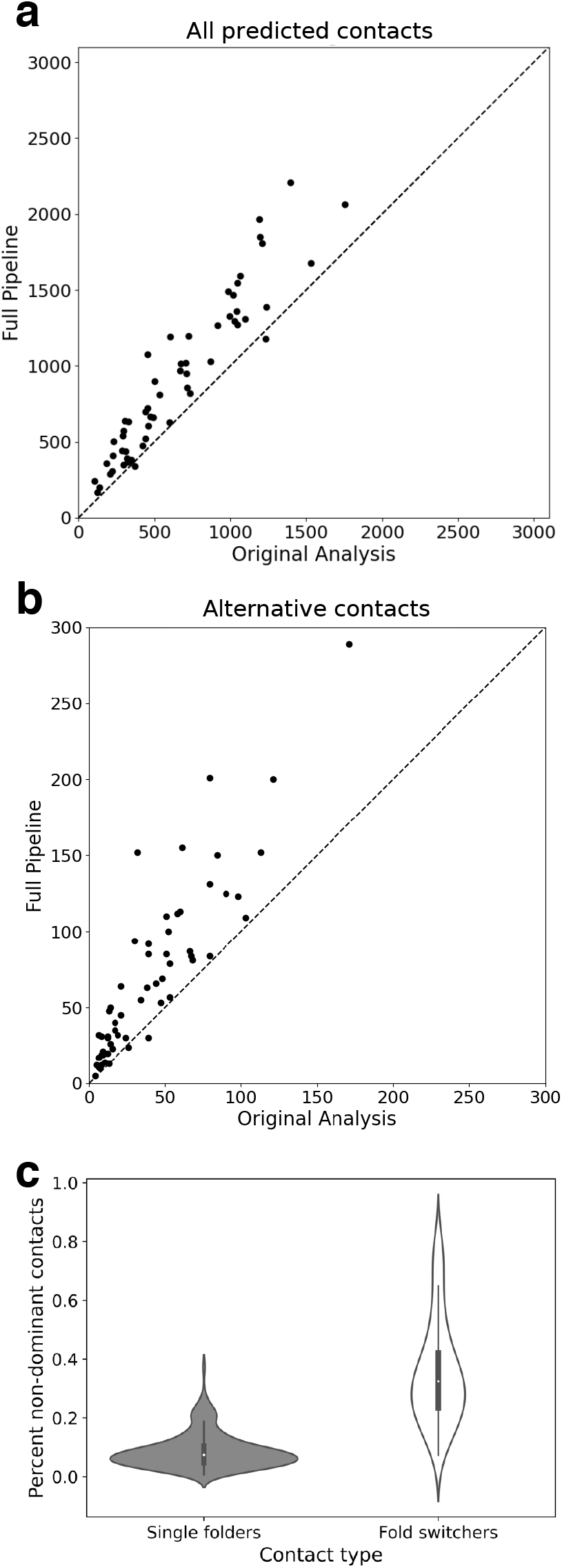
Our workflow amplifies correctly predicted contacts for fold-switching proteins. Amplification is observed for 56/58 predicted contacts (**a**) and especially for contacts uniquely corresponding to the alternative fold (**b**). Identity lines in both plots are dashed lines. (**c**) Noise generated by our pipeline was significantly reduced by density-based filtering. Violin plots show the distributions of %non-dominant contacts for single-fold and fold-switching proteins. The left and right distributions were generated from n=181 and n=58 datapoints, respectively. Inner bold black boxes span the interquartile ranges (IQRs) of each distribution (first quartile, Q1 through third quartile, Q3); medians of each distribution are white dots, lower line (whisker) is the lowest datum above Q1-1.5*IQR; upper line (whisker) is the highest datum below Q3 + 1.5*IQR.

Statistical analysis confirmed that the additional coevolutionary contacts identified by our approach are products of evolution rather than chance. Specifically, the likelihood of generating the additional correct contacts–with concomitant noise–was very low for all 58 proteins, with p-values ranging from 10^−6^ to 0 (one-tailed hypergeometric test, **Extended Data Table 1**). These low p-values demonstrate that the dual-fold coevolutionary signatures identified by GREMLIN and MSA Transformer were almost certainly not generated by chance. Instead, these statistically significant dual-fold signatures suggest that evolution has selected for protein sequences that assume two distinct folds. Furthermore, the distribution of non-dominant contacts for single-fold proteins was significantly lower than for fold switchers (p < 1.1*10^−94^ Epps-Singleton test, **Figure 3c, Extended Data Table 3**). Because our approach especially enriches the number of contacts corresponding to the alternative conformation (**Figure 3b**), these results demonstrate that evolution has selected for protein sequences that assume two distinct folds.

#### Prediction of two folds from one sequence

Widespread dual-fold coevolution opens the possibility of predicting both conformations of a fold-switching protein from its sequence. We tested this possibility on a NusG Variant with low sequence identity (≤29%) to homologs with experimentally determined three-dimensional structures. NusG proteins are the only transcription factors known to be conserved in all kingdoms of life^50^. Unlike most NusGs with atomic level structures, whose C-terminal domains (CTDs) assume a β-roll fold^17,51-54^, this Variant’s CTD switches from an α-helical ground state to a β-roll^7^, much like its homolog, RfaH^48^. Nevertheless, AlphaFold2 consistently predicts that the CTD of this Variant assumes a β-roll fold only (**Figure 4a, Extended Data Figure 3**). This prediction corroborates the observations discussed previously: all NusG CTDs are expected to assume β-roll folds (dominant conformation), though a subpopulation can also assume α-helical folds (alternative conformation). To test whether the coevolutionary signal of the β-roll fold might be masking a weaker α-helical signature, we examined the coevolved amino acid pairs identified by our approach. Twenty-one amino acid positions in the CTD only formed coevolved pairs corresponding to the β-roll fold, while positions exclusively forming coevolved pairs corresponding to the α-helical fold numbered only four (**Figure 4a**). To weaken the coevolutionary signal corresponding to the β-roll fold, we changed all 21 positions in the MSA to alanine, the mutation of choice for perturbing structure^55^, except for the sequence of the Variant (**Figure 4b**). From this modified MSA, AlphaFold2 predicted a ground state α-helical structure (**Figure 4b**). The secondary structures of both CTDs have high prediction confidences (pLDDT scores), except for the most C-terminal helix in the α-hairpin conformation (**Extended Data Figure 4**).

**Figure 4.**
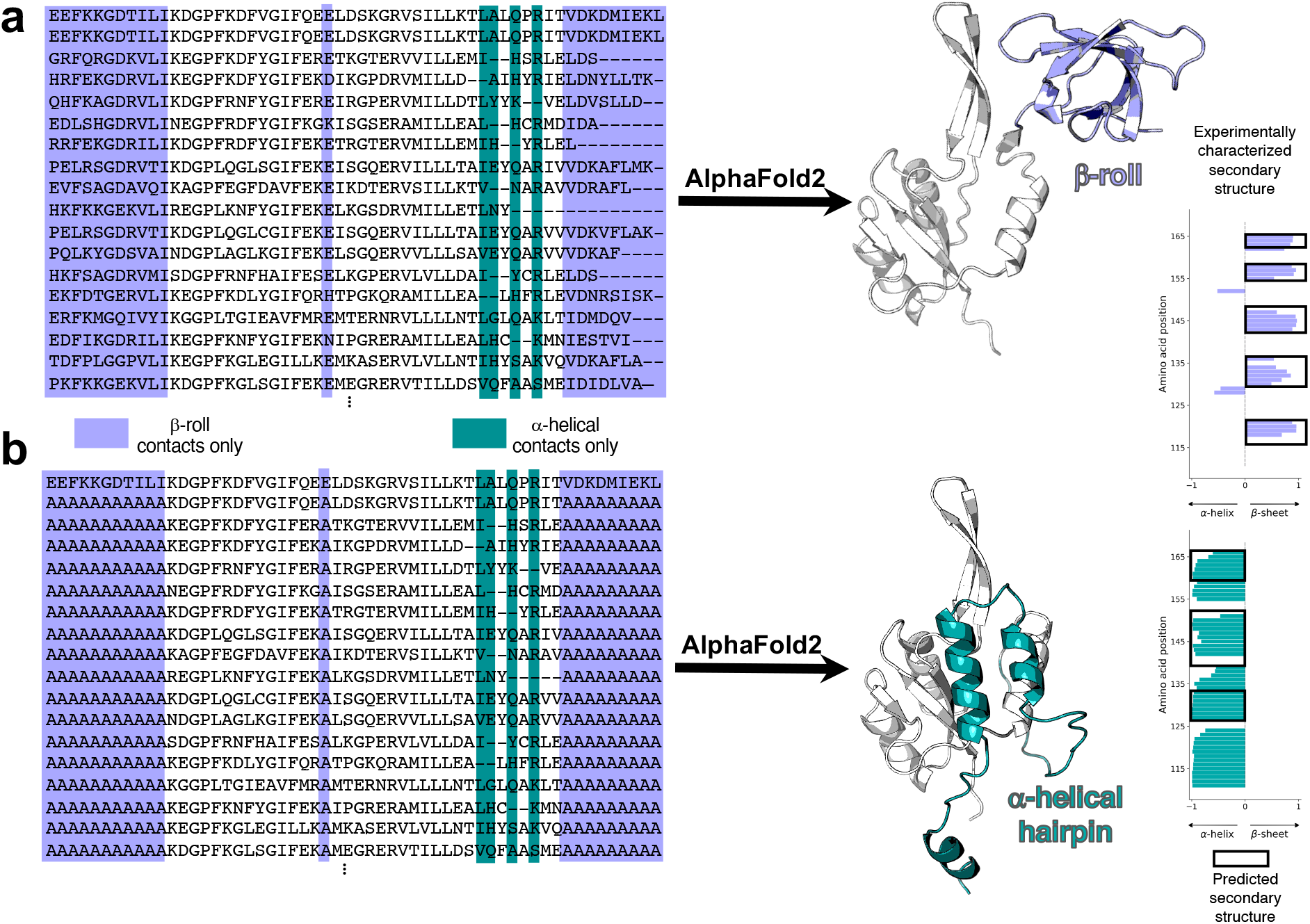
AlphaFold2 successfully predicts two conformations of a candidate sequence without experimentally determined structures. (**a**). A NusG N-terminal (NGN) fold (light gray) and a C-terminal β-roll fold (lavender) are predicted from a deep input MSA (region corresponding to the CTD shown). Predicted β-sheets in the C-terminal domain agree closely with secondary structures predicted from nuclear magnetic resonance experiments (black boxes surrounding lavender bars). (**b**). A NusG N-terminal (NGN) fold (light gray) and a C-terminal α-helical hairpin fold (teal) are predicted from a modified input MSA in which columns predicted to form only β-roll contacts are changed to alanine. Predicted α-helices in the C-terminal domain agree with secondary structures predicted from nuclear magnetic resonance experiments (black boxes surrounding teal bars). Protein structures were generated with PyMOL^46^.

Both predicted conformations are consistent with amino-acid-specific secondary structure predictions calculated from nuclear magnetic resonance assignments^7^ (**Figure 4**). Furthermore, without suppressing the strong β-roll coevolutionary signature, AlphaFold2 consistently predicted the β-sheet fold regardless of input MSAs and use or absence of templates. The α-helical CTD conformation was also missed by RoseTTAfold^3^ and RGN2^56^, an MSA-independent deep learning method that outperforms AlphaFold2 on orphan protein sequences (**Extended Data Figure 3**). Together, these results demonstrate that the coevolved contacts identified by our approach facilitated predictions of two folds from one sequence.

## Discussion

Although globular proteins are generally observed to assume single unique folds, an increasing number can switch between distinct sets of stable secondary and tertiary structure. These fold-switching proteins facilitate cancer progression^57^, foster SARS-CoV-2 pathogenesis^58^, fight microbial infection^37^, and more^20^.

By running well-developed coevolutionary analysis methods^35,42,45^ on many sets of protein superfamilies and subfamilies, we identified statistically significant coevolutionary signals corresponding to two folds of 58 diverse fold-switching proteins. This widespread selection indicates that fold switching (1) confers evolutionary advantage and (2) is a fundamental biological mechanism. These results, coupled with the difficulties associated with experimentally characterizing fold switchers^2,14^, suggest that fold-switching proteins may be more naturally abundant than currently realized. Accordingly, recent experimentally confirmed predictions suggest that over 3500 proteins in the NusG transcription factor family of ∼15,500 proteins switch folds^7^. Furthermore, since subfamily MSAs have also been used to infer other protein properties^59,60^, our approach might successfully extend beyond fold switchers to other forms of structural heterogeneity, such as allostery, which previous coevolutionary approaches have predicted with some success^61,62^.

The observed prevalence and biological relevance of fold-switching proteins underscore the need to develop computational methods that reliably predict more. Although state-of-the-art predictive algorithms have revolutionized protein structure prediction^3,5,8^, they systematically fail to predict protein fold switching^6,7^. Nevertheless, we show here that AlphaFold2 can be biased to predict two folds from one amino acid sequence. The key to this approach was suppressing the strong coevolutionary signature of the β-roll fold, allowing AlphaFold2 to detect weaker α-helical signals. This approach may extend to other fold-switching proteins^55^, though a systematic analysis has not yet been performed. Furthermore, a recent preprint reported that AlphaFold2 could be guided to predict both structures of 3/6 known fold-switching proteins by clustering their sequences into subfamilies^63^.

Our findings lay the groundwork for a more functionally complete picture of the proteome by capturing dual-fold coevolutionary signatures of fold-switching proteins from their genomic sequences. Still, further technical advances are needed to reliably predict protein fold switching. First, dual-fold contacts must be distinguished from noise or true contacts arising from other phenomena, such as multimerization. Second, dual-fold contacts must be correctly separated into their two respective folds without the knowledge of both conformations, on which we rely here. Third, robust dual-fold contacts must be predicted reliably. Our approach works only on sequences for which deep MSAs can be generated. As a result, fold switching could not be predicted in nearly 50% of the sequences in our initial dataset. Furthermore, it is uncertain how many sets of the dual-fold contacts we generated are complete enough to robustly predict two distinct folds from single amino acid sequences. Nevertheless, the rapid growth of diverse protein sequence^64^, recent advances in deep learning^65,66^, and increasingly accessible computational resources leave us optimistic that these challenges will be overcome.

## Materials and Methods

### MSA generation

Fold-switching protein sequences were used as inputs for jackhmmer^67,68^ to generate multiple sequence alignments (MSAs) after searching the Uniref90^64^ release from January 2021. To achieve optimal MSA depths, multiple searches with -incE and -incdomE thresholds set to the same value ranging from 10^−1^ to 10^−250^ were performed in increments of 10^−3^. We then searched for the deepest MSA in this range with a maximum of 60,000 sequences. Each jackhmmer run was iterated until the MSA converged or until 10 iterations had occurred.

### MSA preparation

To generate subfamily MSAs, distantly related sequences were pruned from deep superfamily MSAs using hhfilter^69^. This software filters alignments by QID, pairwise sequence identity between the query sequence used to generate the MSA and each subsequent sequence within it. Subfamily MSAs of varying depths were generated with QID thresholds ranging from 1% to 50% in increments of 1%. All MSAs—both superfamily and subfamily—were prepared for coevolutionary analysis by removing any sequences with >25% gaps and then filtering any columns with >75% gaps.

### Coevolutionary Analysis

Prepared MSAs from each protein family were used as separate inputs into both GREMLIN^41,42^ and MSA transformer^35^, each run with default parameters. Typically, the number of coevolved amino acid pairs retained from each run from both programs is 3*L*/2^45,70^, where *L* is length of the target protein. Here, 2*L* pairs are retained for the deepest MSA because we expect more coevolved contacts to arise from fold-switching sequences with two conformations. The number of contacts retained from subfamilies decreases linearly by the number of sequences in the subfamily MSA normalized by the number of sequences in the deepest MSA to a minimum value of 3*L*/2 for the shallowest input MSA from each protein family. The *N* contacts with the highest APC scores for each GREMLIN run and the *M* contacts with the highest Z-scores from each MSA Transformer run were retained, where *N and M* ∈ [3*L*/2,2*L*]. Predicted contacts from each MSA were combined and duplicates were removed.

### Noise filtering

All predicted contacts generated from the original MSA and the sub-family MSAs were superimposed onto a single contact map. These predictions were clustered using a density-based algorithm (DBSCAN)^49^ that efficiently identifies structure in datasets with arbitrarily shaped clusters. The main criteria for defining whether a point belongs to a cluster is how many other points are close. The EPS parameter defines a radial distance from a core point and points within that radius are clustered. All points included in the cluster were then used as new core points to search for additional points within the EPS. Clusters were iteratively built in this way until the entire dataset is clustered. The minimum number of points to define a cluster in this work is 3. The sparsest points in the dataset were then defined as noise and eliminated from the dataset to produce the final, densest set of filtered predictions. The EPS value is optimized for each set of superimposed contacts using a receiver operating characteristic (ROC) curve, where the optimal value’s first derivative > 1, corresponding to more true positives gained by increasing the EPS value, but the successive value’s first derivative <1, corresponding to more false positives gained by further increasing the EPS value. True positives are defined as being within +/-1 residue of crystallographic contacts. However, EPS values could not be so stringent that fewer contacts were returned than from the original run on deep MSAs.

### Statistical tests

p-values were calculated using the one-tailed hypergeometric test (also known as Fisher’s exact test) to evaluate the significance of the additional structural information obtained from the subfamily alignments, as described by:

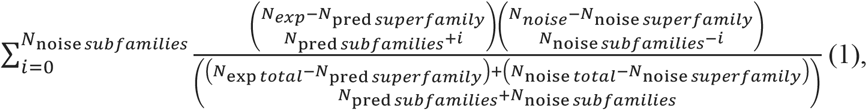

where *N*_*exp*_ is the total number of unique experimentally determined contacts from both conformations of a fold-switching protein, *N*_pred *superfamily*_ is the number of unique contacts correctly predicted by GREMLIN and MSA transformer on the superfamily MSA only, *N*_pred *superfamilies*._ is the number of unique contacts predicted by GREMLIN and MSA transformer on all subfamily MSAs excluding those also predicted from the superfamily, *N*_*noise*_ is *L*^2^ − *N*_*exp*_, where *L* is the maximum sequence length of an experimentally determined structure, *N*_noise *superfamily*_ is the number of unique contacts incorrectly predicted by GREMLIN or MSA transformer on the superfamily MSA only, and *N*_noise *superfamilies*_ is the number of unique contacts incorrectly predicted by GREMLIN or MSA transformer on all subfamily MSAs, excluding those also predicted from the superfamily.

### Structure predictions

Structure predictions of Variant 5 were performed by AlphaFold2.1.2 both with templates deposited in the PDB by 4/20/22 and without templates and both with MSAs generated from the standard pipeline (Uniref90^71^, MGnify^72^, and MMseqs2^73^ (BFD clust)) and the shallowest MSA generated from our approach. In all four runs, only the β-roll fold was predicted in the five top-scoring models (**Figure S3**). The α-helical fold was predicted by modifying MSA columns that our pipeline predicted to form only β-roll contacts. These columns corresponded to amino acids in the 100-168 range, and the deepest MSA generated by our approach was modified by mutating these columns to alanine. AlphaFold2.1.2 was run on this modified MSA without templates. The standard RoseTTAfold pipeline (https://robetta.bakerlab.org) was used to predict three-dimensional structures of Variant 5 both with the standard MSA generation protocol and the shallowest MSA generated from our pipeline.

Finally, the RGN2 Colab notebook (https://colab.research.google.com/github/aqlaboratory/rgn2/blob/master/rgn2_prediction.ipynb) was run on the sequence of Variant 5 with standard parameters. The sequence of Variant 5 is: MESFLNWYLIYTKVKKEDYLEQLLTEAGLEVLNPKIKKTKTVRNKKKEVIDPLFPCYLFVKADLNVHLRIISYTQGIRRLVGGSNPTIVPIEIIDTIKSRMVDGFIDTKSEEFKKGDTILIKDGPFKDFVGIFQEELDSKGRVSILLKTLALQPRITVDKDMIEKLHN. Experimentally determined secondary structures were taken from^7^.

## Acknowledgments

We thank Loren Looger, Carolyn Ott, Yuri Wolf, Nash Rochman, Robert Best, Andy LiWang, Eugene Koonin, Devlina Chakravarty, and Danielle and Jean Thierry-Mieg for helpful discussions. This work utilized resources from the NIH HPS Biowulf cluster (http://hpc.nih.gov), and it was supported by the Intramural Research Program of the National Library of Medicine, National Institutes of Health.

