## Extended Data for "Evolutionary selection of proteins with two folds"

*Table S1 (attached separately): Structural information and pipeline output for 54 fold-switching proteins. PDBs A and B (PDB ID_Chain) were used to calculate experimentally determined contacts for a given protein; sequences of blue codes were used queries for generating deep MSAs. Name is the common name of each protein. MSA Depth is the number of sequences in the deepest MSA prepared for GREMLIN and MSA transformer and Neff is the number of effective sequences in the deepest MSA. L is the number of amino acids in the query sequence. GMN_QID and MSATR_QID are the number of subfamily alignments made from the original MSA deep enough to run each algorithm. P-values are calculated using the one-tailed hypergeometric test and represent the significance of the additional structural information obtained from the subfamily alignments.*

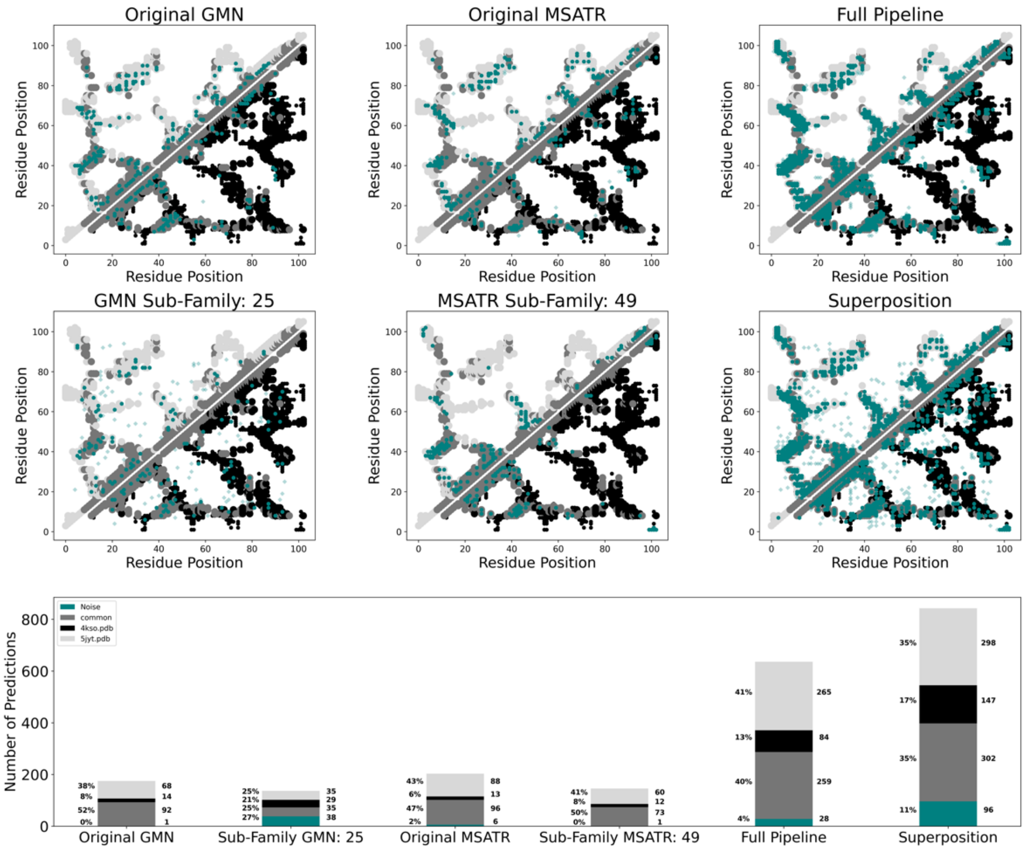


*Figures S1: Our approach enhances predictions of coevolved amino acid pairs: example 1 of 10. All six upper panels show predicted contacts referenced against contacts from two experimentally determined structures of KaiB with PDB codes 5JYT (dominant conformation) and 4KSO (alternative conformation). Color schemes of these six panels follow: experimentally determined contacts unique to the dominant conformation (light gray), alternative conformation (black), common to both conformations (dark gray), interchain contacts from homomers (small circles); predicted contacts (teal) corresponding to experimentally determined structures (opaque circles), not corresponding to experimentally determined structures (noise, translucent diamonds). Panels, top-to-bottom, left-to-right show the experimentally determined contacts underlaying: GREMLIN predictions from the original MSA (Original GMN) and the shallowest subfamily MSA (GMN subfamily, 25 indicates minimum %sequence identity to query); MSA Transformer predictions from the original MSA (Original MSATR) and the shallowest subfamily MSA (MSATR subfamily, 49 indicates minimum %sequence identity to query); predictions from our approach after noise filtering (Full Pipeline) and before noise filtering (Superposition). Predicted contacts from each panel are tabulated by overlap with experimentally determined contacts in the bar graph: dominant (light gray); alternative (black); common (dark gray); noise (teal). Predicted contacts from GREMLIN and MSA Transformer are often redundant; signal increases from our approach are shown in Figure 3.*

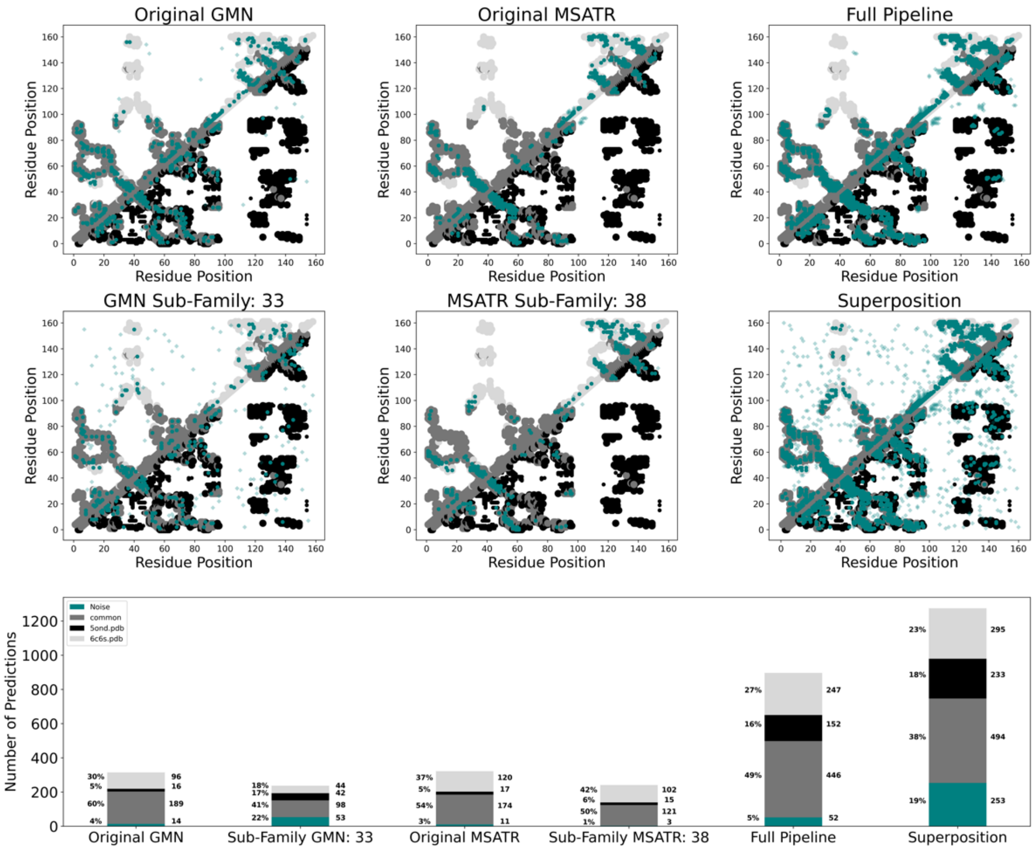


*Figures S1: Our approach enhances predictions of coevolved amino acid pairs: example 2 of 10. All six upper panels show predicted contacts referenced against contacts from two experimentally determined structures of RfaH with PDB codes 6C6S (dominant conformation) and 5OND (alternative conformation). Color schemes of these six panels follow: experimentally determined contacts unique to the dominant conformation (light gray), alternative conformation (black), common to both conformations (dark gray), interchain contacts from homomers (small circles); predicted contacts (teal) corresponding to experimentally determined structures (opaque circles), not corresponding to experimentally determined structures (noise, translucent diamonds). Panels, top-to-bottom, left-to-right show the experimentally determined contacts underlaying: GREMLIN predictions from the original MSA (Original GMN) and the shallowest subfamily MSA (GMN subfamily, 33 indicates minimum %sequence identity to query); MSA Transformer predictions from the original MSA (Original MSATR) and the shallowest subfamily MSA (MSATR subfamily, 38 indicates minimum %sequence identity to query); predictions from our approach after noise filtering (Full Pipeline) and before noise filtering (Superposition). Predicted contacts from each panel are tabulated by overlap with experimentally determined contacts in the bar graph: dominant (light gray); alternative (black); common (dark gray); noise (teal). Predicted contacts from GREMLIN and MSA Transformer are often redundant; signal increases from our approach are shown in Figure 3*

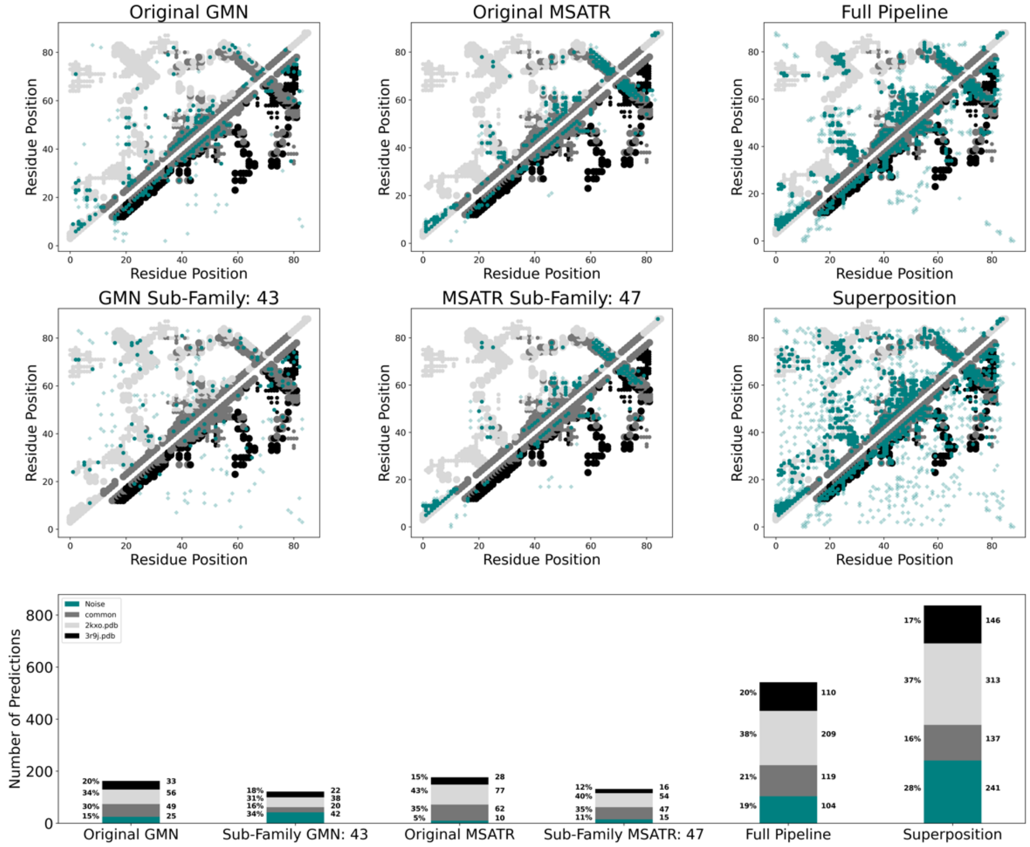


*Figures S1: Our approach enhances predictions of coevolved amino acid pairs: example 3 of 10. All six upper panels show predicted contacts referenced against contacts from two experimentally determined structures of MinE with PDB codes 2KXO (dominant conformation) and 3R9J (alternative conformation). Color schemes of these six panels follow: experimentally determined contacts unique to the dominant conformation (light gray), alternative conformation (black), common to both conformations (dark gray), interchain contacts from homomers (small circles); predicted contacts (teal) corresponding to experimentally determined structures (opaque circles), not corresponding to experimentally determined structures (noise, translucent diamonds). Panels, top-to-bottom, left-to-right show the experimentally determined contacts underlaying: GREMLIN predictions from the original MSA (Original GMN) and the shallowest subfamily MSA (GMN subfamily, 43 indicates minimum %sequence identity to query); MSA Transformer predictions from the original MSA (Original MSATR) and the shallowest subfamily MSA (MSATR subfamily, 47 indicates minimum %sequence identity to query); predictions from our approach after noise filtering (Full Pipeline) and before noise filtering (Superposition). Predicted contacts from each panel are tabulated by overlap with experimentally determined contacts in the bar graph: dominant (light gray); alternative (black); common (dark gray); noise (teal). Predicted contacts from GREMLIN and MSA Transformer are often redundant; signal increases from our approach are shown in Figure 3.*

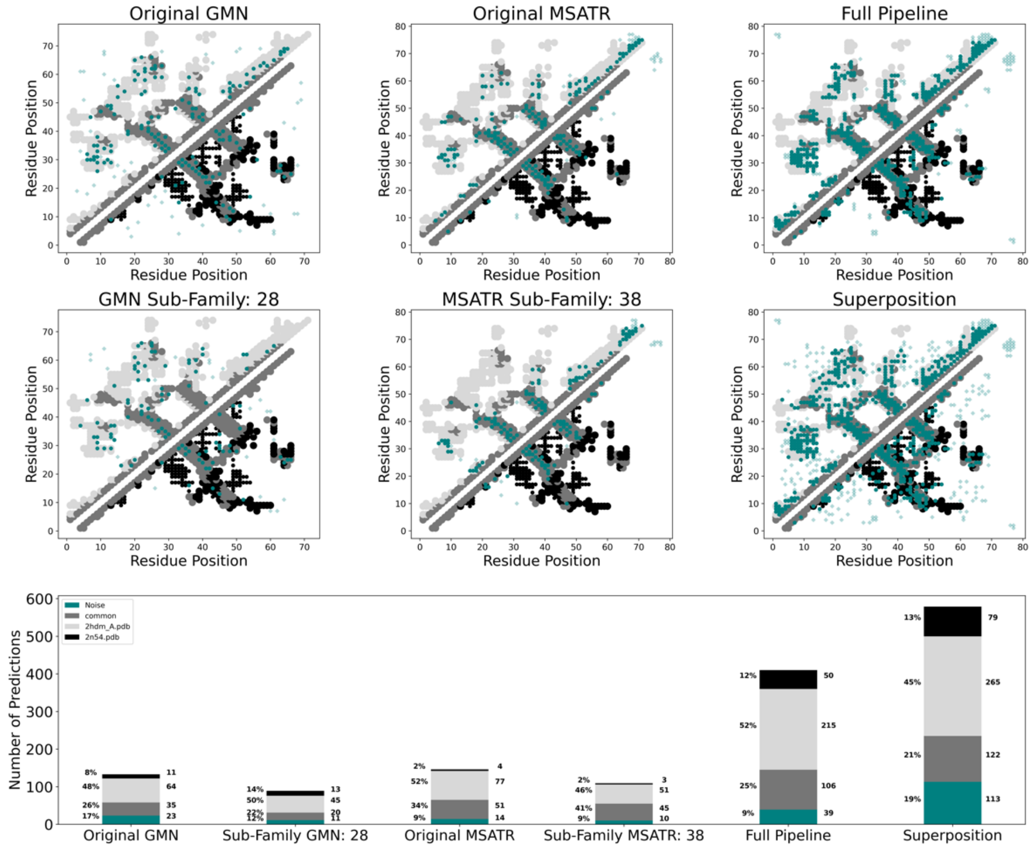


*Figures S1: Our approach enhances predictions of coevolved amino acid pairs: example 4 of 10. All six upper panels show predicted contacts referenced against contacts from two experimentally determined structures of XCL1 with PDB codes 2HDM (dominant conformation) and 2N54 (alternative conformation). Color schemes of these six panels follow: experimentally determined contacts unique to the dominant conformation (light gray), alternative conformation (black), common to both conformations (dark gray), interchain contacts from homomers (small circles); predicted contacts (teal) corresponding to experimentally determined structures (opaque circles), not corresponding to experimentally determined structures (noise, translucent diamonds). Panels, top-to-bottom, left-to-right show the experimentally determined contacts underlaying: GREMLIN predictions from the original MSA (Original GMN) and the shallowest subfamily MSA (GMN subfamily, 28 indicates minimum %sequence identity to query); MSA Transformer predictions from the original MSA (Original MSATR) and the shallowest subfamily MSA (MSATR subfamily, 38 indicates minimum %sequence identity to query); predictions from our approach after noise filtering (Full Pipeline) and before noise filtering (Superposition). Predicted contacts from each panel are tabulated by overlap with experimentally determined contacts in the bar graph: dominant (light gray); alternative (black); common (dark gray); noise (teal). Predicted contacts from GREMLIN and MSA Transformer are often redundant; signal increases from our approach are shown in Figure 3.*

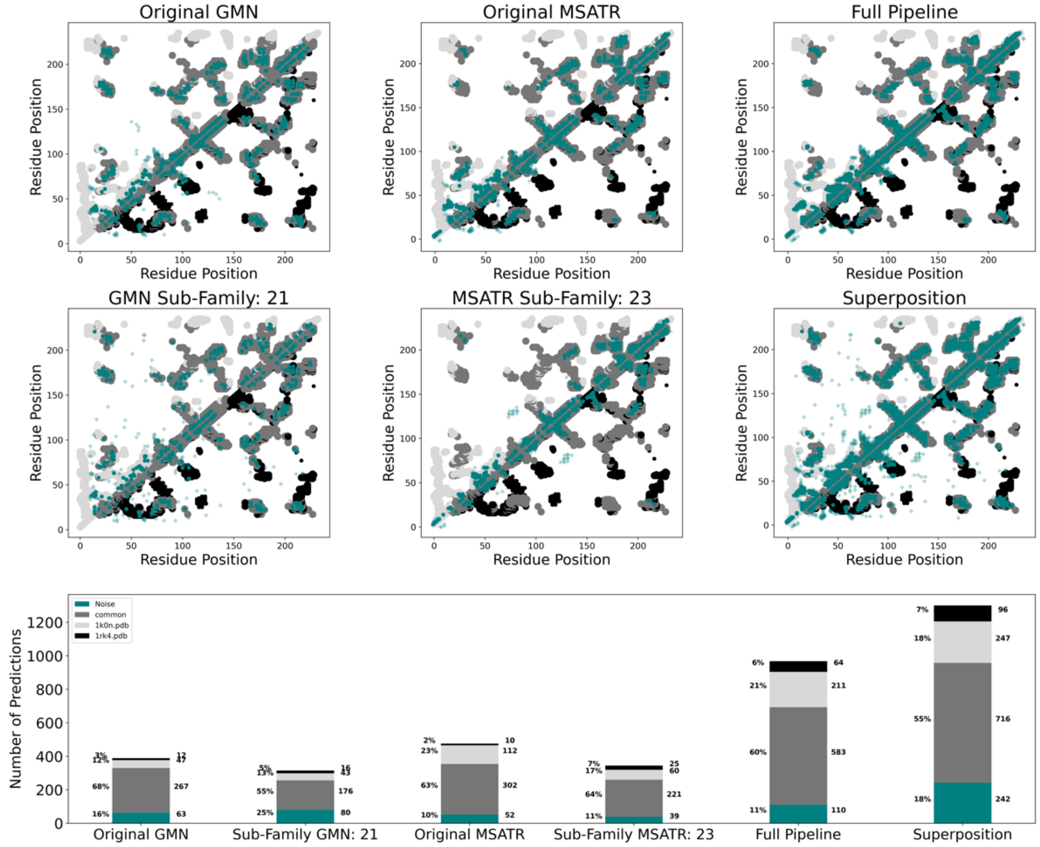


*Figures S1: Our approach enhances predictions of coevolved amino acid pairs: example 5 of 10. All six upper panels show predicted contacts referenced against contacts from two experimentally determined structures of CLIC1 with PDB codes 1K0N (dominant conformation) and 1RK4 (alternative conformation). Color schemes of these six panels follow: experimentally determined contacts unique to the dominant conformation (light gray), alternative conformation (black), common to both conformations (dark gray), interchain contacts from homomers (small circles); predicted contacts (teal) corresponding to experimentally determined structures (opaque circles), not corresponding to experimentally determined structures (noise, translucent diamonds). Panels, top-to-bottom, left-to-right show the experimentally determined contacts underlaying: GREMLIN predictions from the original MSA (Original GMN) and the shallowest subfamily MSA (GMN subfamily, 21 indicates minimum %sequence identity to query); MSA Transformer predictions from the original MSA (Original MSATR) and the shallowest subfamily MSA (MSATR subfamily, 23 indicates minimum %sequence identity to query); predictions from our approach after noise filtering (Full Pipeline) and before noise filtering (Superposition). Predicted contacts from each panel are tabulated by overlap with experimentally determined contacts in the bar graph: dominant (light gray); alternative (black); common (dark gray); noise (teal). Predicted contacts from GREMLIN and MSA Transformer are often redundant; signal increases from our approach are shown in Figure 3.*

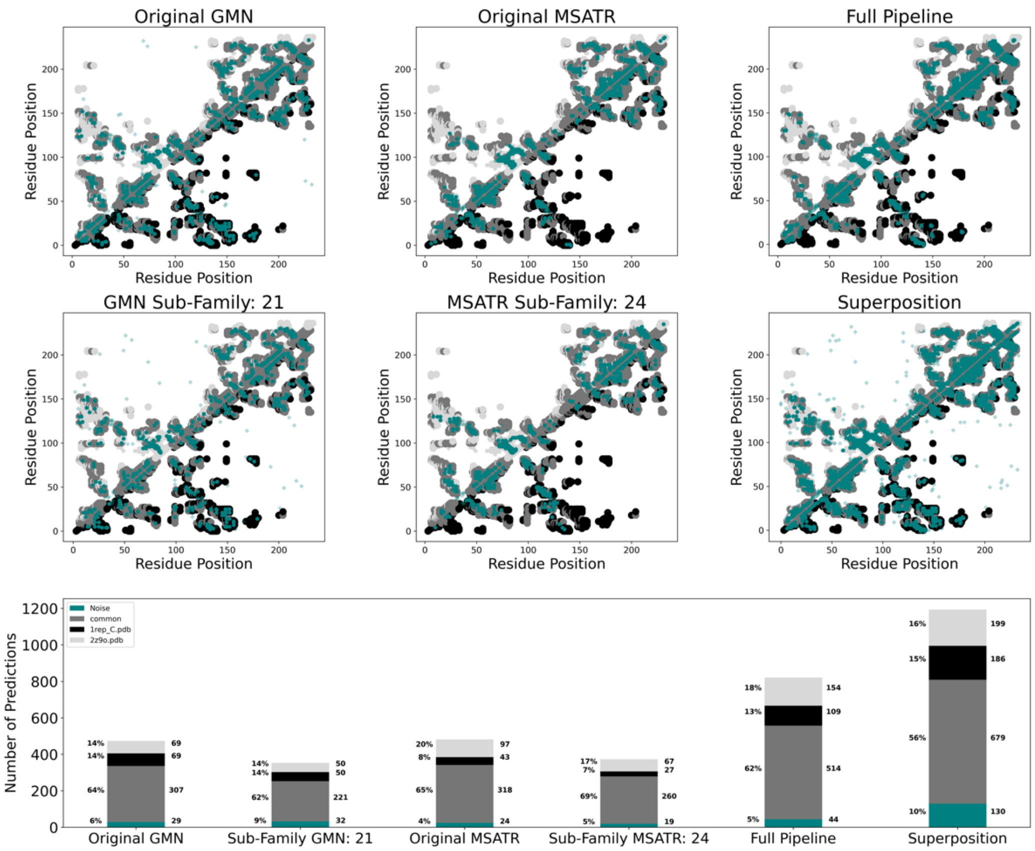


*Figures S1: Our approach enhances predictions of coevolved amino acid pairs: example 6 of 10. All six upper panels show predicted contacts referenced against contacts from two experimentally determined structures of RepE with PDB codes 2Z9O (dominant conformation) and 1REP (alternative conformation). Color schemes of these six panels follow: experimentally determined contacts unique to the dominant conformation (light gray), alternative conformation (black), common to both conformations (dark gray), interchain contacts from homomers (small circles); predicted contacts (teal) corresponding to experimentally determined structures (opaque circles), not corresponding to experimentally determined structures (noise, translucent diamonds). Panels, top-to-bottom, left-to-right show the experimentally determined contacts underlaying: GREMLIN predictions from the original MSA (Original GMN) and the shallowest subfamily MSA (GMN subfamily, 21 indicates minimum %sequence identity to query); MSA Transformer predictions from the original MSA (Original MSATR) and the shallowest subfamily MSA (MSATR subfamily, 24 indicates minimum %sequence identity to query); predictions from our approach after noise filtering (Full Pipeline) and before noise filtering (Superposition). Predicted contacts from each panel are tabulated by overlap with experimentally determined contacts in the bar graph: dominant (light gray); alternative (black); common (dark gray); noise (teal). Predicted contacts from GREMLIN and MSA Transformer are often redundant; signal increases from our approach are shown in Figure 3.*

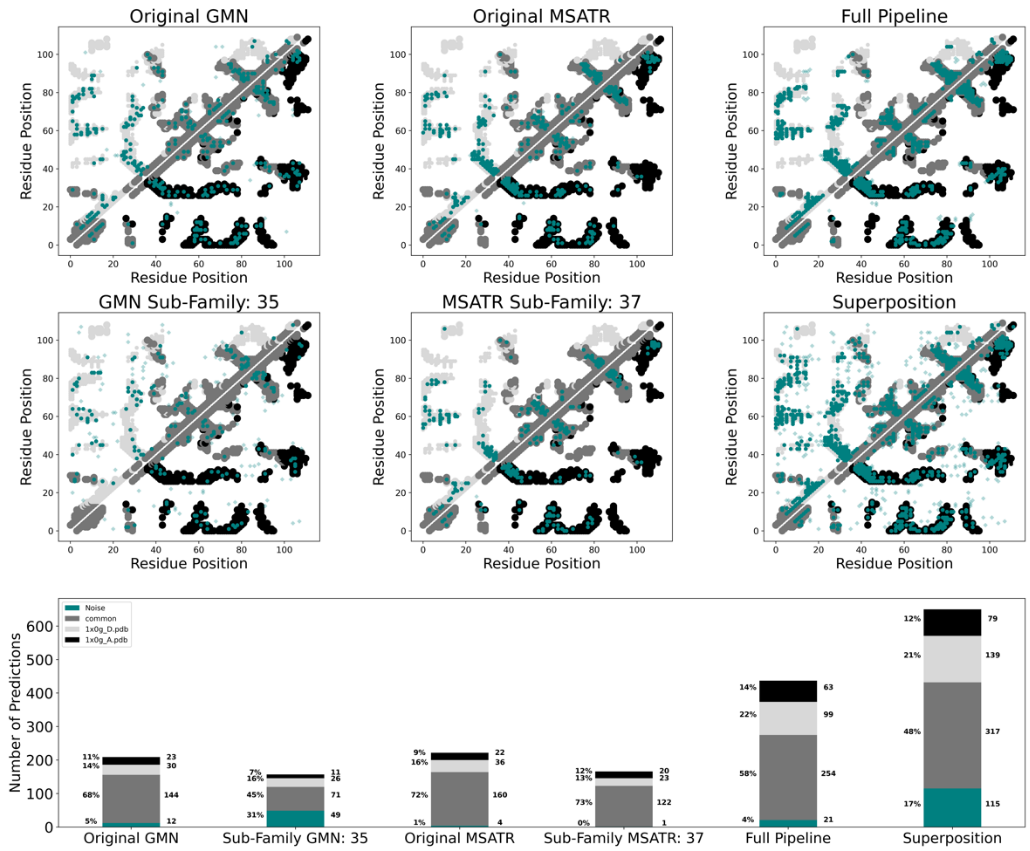


*Figures S1: Our approach enhances predictions of coevolved amino acid pairs: example 7 of 10. All six upper panels show predicted contacts referenced against contacts from two experimentally determined structures of IscA with PDB codes 1X0G chain D (dominant conformation) and 1X0G chain A (alternative conformation). Color schemes of these six panels follow: experimentally determined contacts unique to the dominant conformation (light gray), alternative conformation (black), common to both conformations (dark gray), interchain contacts from homomers (small circles); predicted contacts (teal) corresponding to experimentally determined structures (opaque circles), not corresponding to experimentally determined structures (noise, translucent diamonds). Panels, top-to-bottom, left-to-right show the experimentally determined contacts underlaying: GREMLIN predictions from the original MSA (Original GMN) and the shallowest subfamily MSA (GMN subfamily, 35 indicates minimum %sequence identity to query); MSA Transformer predictions from the original MSA (Original MSATR) and the shallowest subfamily MSA (MSATR subfamily, 37 indicates minimum %sequence identity to query); predictions from our approach after noise filtering (Full Pipeline) and before noise filtering (Superposition). Predicted contacts from each panel are tabulated by overlap with experimentally determined contacts in the bar graph: dominant (light gray); alternative (black); common (dark gray); noise (teal). Predicted contacts from GREMLIN and MSA Transformer are often redundant; signal increases from our approach are shown in Figure 3.*

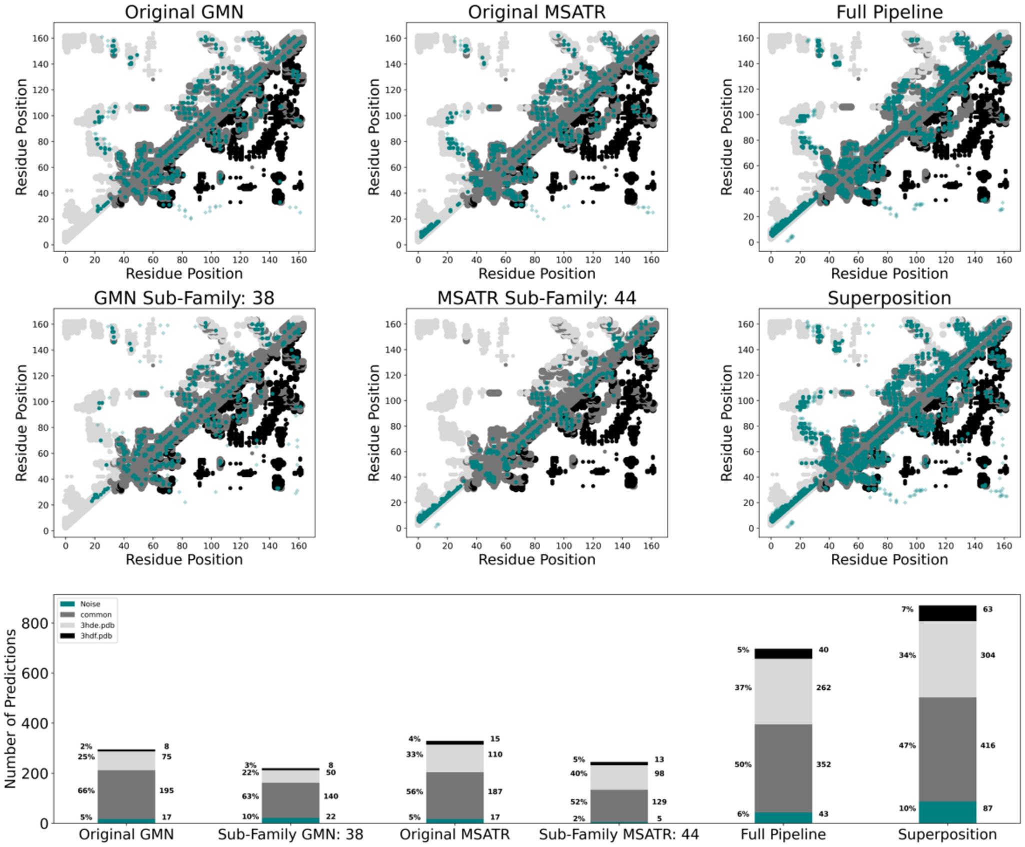


*Figures S1: Our approach enhances predictions of coevolved amino acid pairs: example 8 of 10. All six upper panels show predicted contacts referenced against contacts from two experimentally determined structures of Endolysin with PDB codes 3HDE (dominant conformation) and 3HDF (alternative conformation). Color schemes of these six panels follow: experimentally determined contacts unique to the dominant conformation (light gray), alternative conformation (black), common to both conformations (dark gray), interchain contacts from homomers (small circles); predicted contacts (teal) corresponding to experimentally determined structures (opaque circles), not corresponding to experimentally determined structures (noise, translucent diamonds). Panels, top-to-bottom, left-to-right show the experimentally determined contacts underlaying: GREMLIN predictions from the original MSA (Original GMN) and the shallowest subfamily MSA (GMN subfamily, 38 indicates minimum %sequence identity to query); MSA Transformer predictions from the original MSA (Original MSATR) and the shallowest subfamily MSA (MSATR subfamily, 44 indicates minimum %sequence identity to query); predictions from our approach after noise filtering (Full Pipeline) and before noise filtering (Superposition). Predicted contacts from each panel are tabulated by overlap with experimentally determined contacts in the bar graph: dominant (light gray); alternative (black); common (dark gray); noise (teal). Predicted contacts from GREMLIN and MSA Transformer are often redundant; signal increases from our approach are shown in Figure 3.*

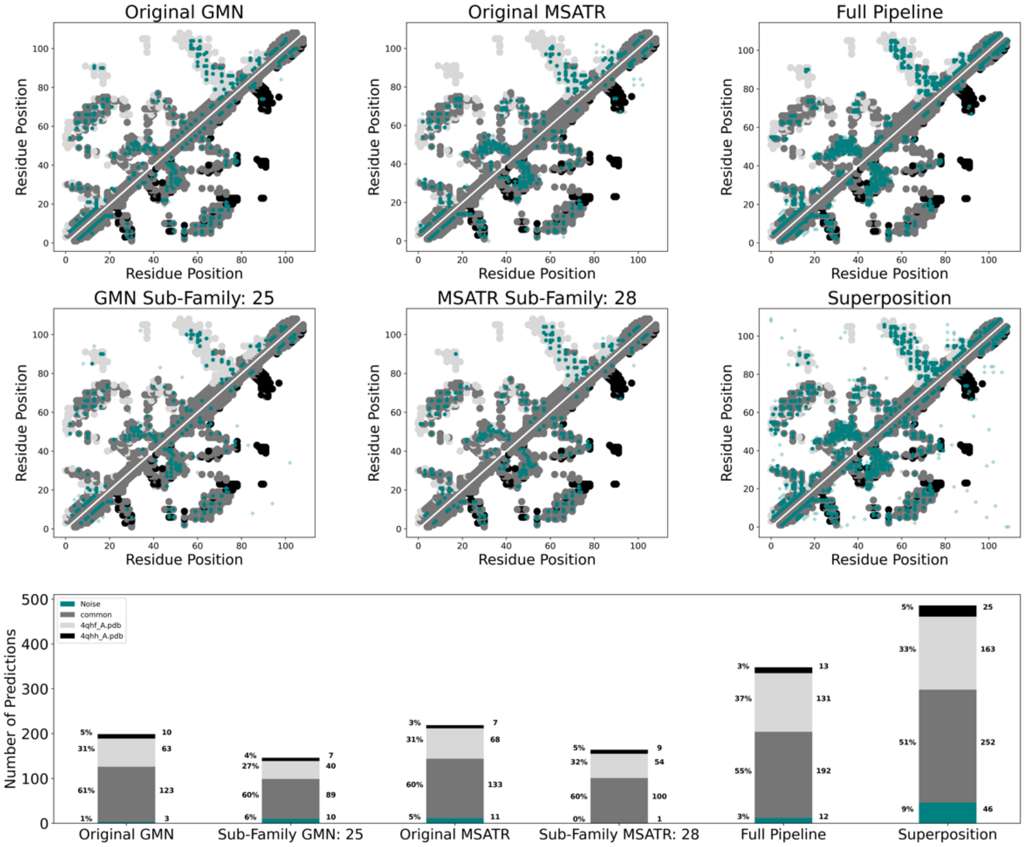


*Figures S1: Our approach enhances predictions of coevolved amino acid pairs: example 9 of 10. All six upper panels show predicted contacts referenced against contacts from two experimentally determined structures of Selecase with PDB codes 4QHF (dominant conformation) and 4QHH (alternative conformation). Color schemes of these six panels follow: experimentally determined contacts unique to the dominant conformation (light gray), alternative conformation (black), common to both conformations (dark gray), interchain contacts from homomers (small circles); predicted contacts (teal) corresponding to experimentally determined structures (opaque circles), not corresponding to experimentally determined structures (noise, translucent diamonds). Panels, top-to-bottom, left-to-right show the experimentally determined contacts underlaying: GREMLIN predictions from the original MSA (Original GMN) and the shallowest subfamily MSA (GMN subfamily, 25 indicates minimum %sequence identity to query); MSA Transformer predictions from the original MSA (Original MSATR) and the shallowest subfamily MSA (MSATR subfamily, 28 indicates minimum %sequence identity to query); predictions from our approach after noise filtering (Full Pipeline) and before noise filtering (Superposition). Predicted contacts from each panel are tabulated by overlap with experimentally determined contacts in the bar graph: dominant (light gray); alternative (black); common (dark gray); noise (teal). Predicted contacts from GREMLIN and MSA Transformer are often redundant; signal increases from our approach are shown in Figure 3.*

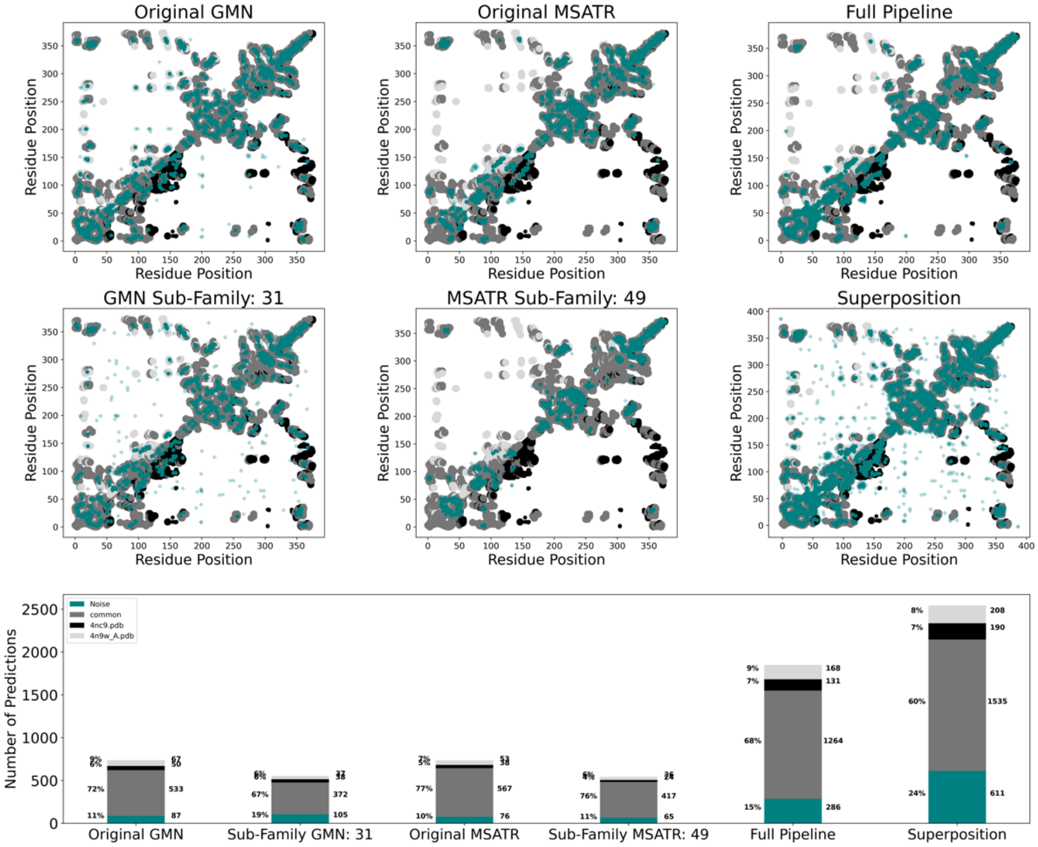


*Figures S1: Our approach enhances predictions of coevolved amino acid pairs: example 10 of 10. All six upper panels show predicted contacts referenced against contacts from two experimentally determined structures of PimA with PDB codes 4N9W (dominant conformation) and 4NC9 (alternative conformation). Color schemes of these six panels follow: experimentally determined contacts unique to the dominant conformation (light gray), alternative conformation (black), common to both conformations (dark gray), interchain contacts from homomers (small circles); predicted contacts (teal) corresponding to experimentally determined structures (opaque circles), not corresponding to experimentally determined structures (noise, translucent diamonds). Panels, top-to-bottom, left-to-right show the experimentally determined contacts underlaying: GREMLIN predictions from the original MSA (Original GMN) and the shallowest subfamily MSA (GMN subfamily, 31 indicates minimum %sequence identity to query); MSA Transformer predictions from the original MSA (Original MSATR) and the shallowest subfamily MSA (MSATR subfamily, 49 indicates minimum %sequence identity to query); predictions from our approach after noise filtering (Full Pipeline)*

*Table S2. Correspondence between experimentally observed and dominant contacts. Conformations are ordered to be experimentally consistent; protein pairs whose predictions do not match experiment are highlighted orange. Conformations equally populated at equilibrium are marked with **. PDB ID format: ID_Chain.*

| Protein Pair | Dominant | Alternative | Protein Pair | Dominant | Alternative |
|---|---|---|---|---|---|
| 1 | 2k42_A | 1cee_B | 30 | 3ejh_A | 3m7p_A |
| 2 | 5fhc_J | 1ebo_E | 31 | 3hde_A | 3hdf_A |
| 3 | 1iyt_A | 2nao_F | 32 | 5et5_A | 3ifa_A |
| 4 | 1jfk_A | 2nxq_B | 33 | 3j7w_B** | 3j7v_G** |
| 5 | 1jti_A | 1ova_A | 34 | 3m1b_F | 3low_A |
| 6 | 1k0n_A | 1rk4_B | 35 | 3uyi_A | 3v0t_A |
| 7 | 1kct_A | 3t1p_A | 36 | 3zwg_N | 4tsy_D |
| 8 | 1qs8_B | 1miq_B | 37 | 4aan_A | 4aal_A |
| 9 | 1mnm_D** | 1mnm_C** | 38 | 4dxt_A | 4dxr_A |
| 10 | 1nqj_B | 1nqd_A | 39 | 4gqc_C | 4gqc_B |
| 11 | 1rep_C | 2z9o_B | 40 | 4jph_B | 5hk5_H |
| 12 | 2h44_A | 1rkp_A | 41 | 4n9w_A | 4nc9_C |
| 13 | 4kso_A | 5jyt_A | 42 | 4o0p_A | 4o01_D |
| 14 | 1uxm_K | 2nam_A | 43 | 4pyi_A | 4pyj_A |
| 15 | 1x0g_D** | 1x0g_A** | 44 | 4uv2_D | 4q79_F |
| 16 | 1xjt_A | 1xju_B | 45 | 4qhf_A | 4qhh_A |
| 17 | 3jv6_A | 1zk9_A | 46 | 4rwn_A | 4rwq_B |
| 18 | 2grm_B | 2axz_A | 47 | 4twa_A | 4ydq_B |
| 19 | 2hdm_A** | 2n54_B** | 48 | 4y0m_J | 4xws_D |
| 20 | 2jmr_A | 4j3o_F | 49 | 4zrb_C | 4zrb_H |
| 21 | 2k0q_A | 2lel_A | 50 | 5b3z_A | 5bmy_A |
| 22 | 2kxo_A | 3rj9_C | 51 | 5c1v_A | 5c1v_B |
| 23 | 2lep_A | 4hdd_A | 52 | 5f5r_B | 5f3k_A |
| 24 | 2mwf_A | 2nnt_A | 53 | 5ond_A | 6c6s_D |
| 25 | 2p3v_A** | 2p3v_D** | 54 | 2c1v_B | 2c1u_C |
| 26 | 2qqj_A | 4qds_A | | | |
| 27 | 2uy7_D | 5flu_E | | | |
| 28 | 2vfx_L** | 3gmh_L** | | | |
| 29 | 4phq_A | 2wcd_X | | | |

*Table S3. List of 181 single-fold proteins analyzed by our pipeline (attached separately). Distributions of non-dominant contacts from these proteins are shown in Figure 3c.*

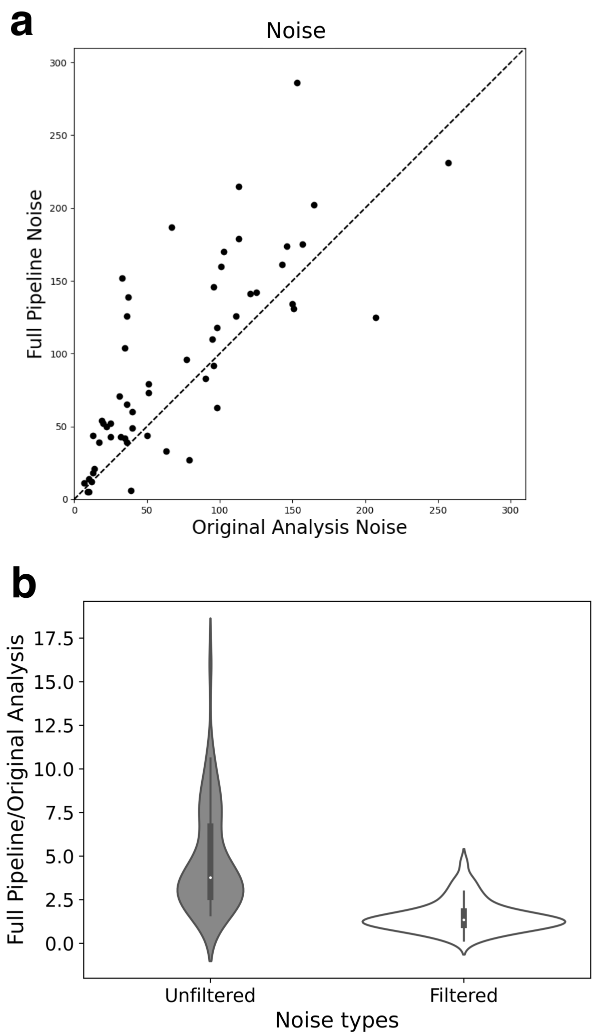


*Figure S2 (a) Comparison of noise from our pipeline with noise from coevolutionary analysis (GREMLIN+MSA Transformer) on deep MSAs only. Mean/median noise increases were 56%/34%, significantly less than the increase in alternative contacts (mean/median 99%/69%, Figure 3b). (b)Violin plots show the distributions of fold increase in noise generated by our pipeline relative to the original analysis before filtering (Unfiltered) and after filtering (Filtered). Both distributions were derived from n=58 datapoints, each corresponding to analysis on one fold-switching protein. Inner bold black boxes span the interquartile ranges (IQRs) of each distribution (first quartile, Q1 through third quartile, Q3); medians of each distribution are white dots, lower line (whisker) is the lowest datum above Q1-1.5*IQR; upper line (whisker) is the highest datum below Q3 + 1.5*IQR.*

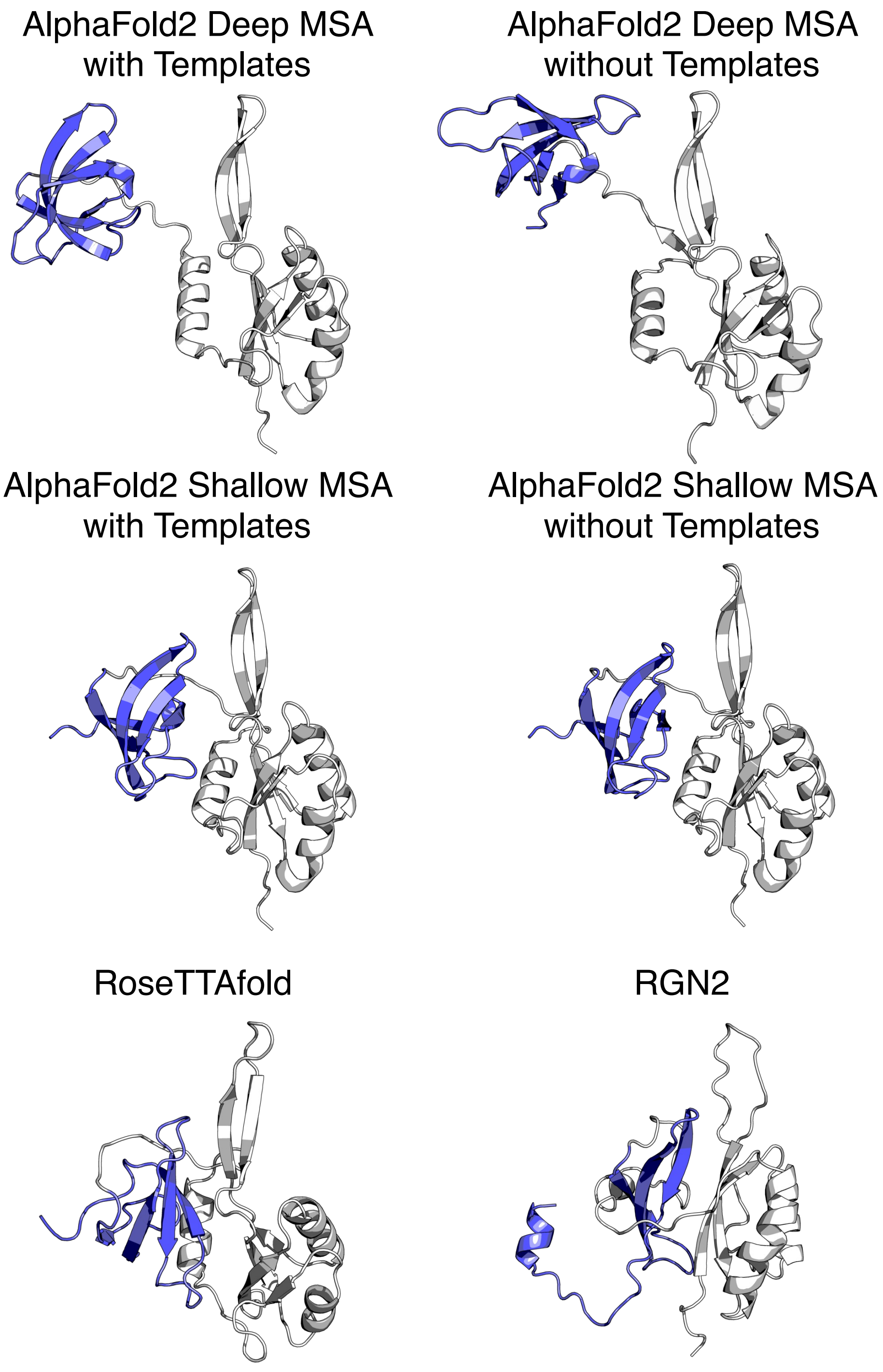


*Figure S3. State-of-the-art methods predict that the C-terminal domain of Variant 5 (blue) assumes conformations mostly composed of $\beta$-sheets. Highest ranked conformers are shown, though the top 5 models from each run all show similar $\beta$-sheet structures. The single-folding N-terminal domain of Variant 5 is shown in white.*

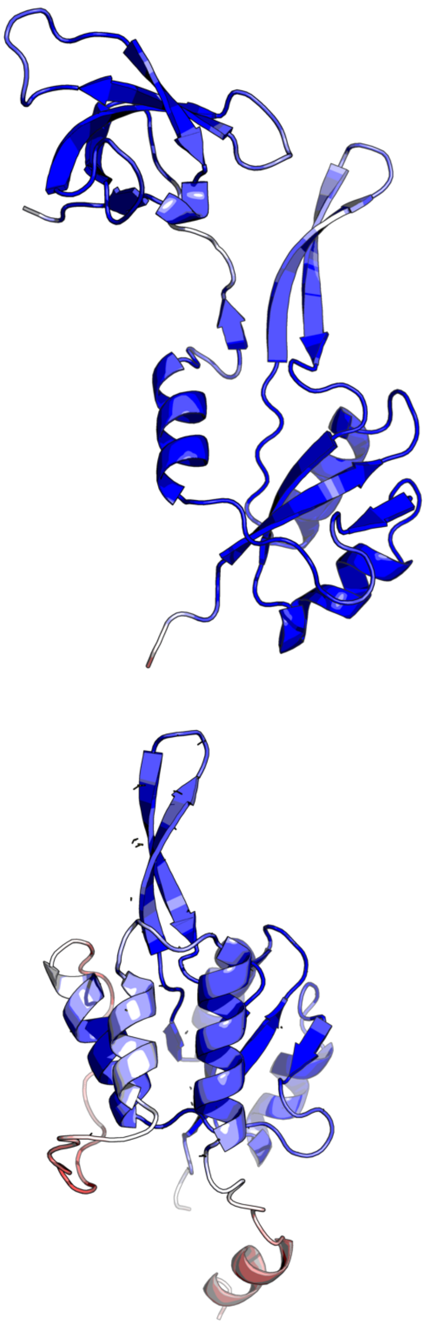

*Figure S4. AlphaFold2 prediction confidences (pLDDT scores) for the $\beta$-roll (above) and $\alpha$-helical (below) conformations of Variant 5. Most confident scores are dark blue; least confident, red, moderate scores are white.*
